# Monooxygenase-dehydrogenase cascade for sustained enzymatic remediation of TMA in salmon protein hydrolysates

**DOI:** 10.1101/2025.06.16.658034

**Authors:** Rasmus Ree, Øivind Larsen, Sushil Gaykawad, Sreerekha S. Ramanand, Antonio García-Moyano, Irina Elena Chiriac, Pål Puntervoll, Gro Elin Kjæreng Bjerga

## Abstract

Fish protein hydrolysates hold great promise as nutraceuticals, yet their application as food ingredients or nutraceuticals is currently limited by their fish-like odor. This odor is mainly due to the presence of trimethylamine (TMA), a volatile biogenic amine resulting from the breakdown of naturally occurring trimethylamine-N-oxide (TMAO) in marine fish. The bacterial trimethylamine monooxygenase mFMO can oxidize TMA into TMAO using molecular oxygen and the cofactor nicotinamide adenine dinucleotide phosphate (NADPH). We have established an enzyme cascade which takes advantage of glucose dehydrogenase to recycle NADPH from NADP^+^, significantly decreasing the cost of the reaction and paving the way for using the enzyme system in fish protein hydrolysates targeted for human consumption. We demonstrate that the dual enzyme system works in an industrially relevant substrate. Salmon protein hydrolysate treated with an mFMO/glucose dehydrogenase cocktail showed a 75% reduction in TMA. A trained sensory panel perceived an improved odor across several parameters, including a reduction in the characteristic TMA smell.

## Introduction

The Norwegian aquaculture and fisheries industry produced more than 3.2 million tons of seafood in 2023 (1). Of this, over 1 million tons were byproducts, defined as any product which is not the main product produced from a raw material, such as heads, frames and backbones. Fish protein hydrolysis is a way of utilizing such byproducts through controlled proteolysis, often using subtilisins (2). The product, fish protein hydrolysates (FPH) is a mix of amino acids and small peptides which are suitable for human consumption. It is an excellent protein source, with protein content varying depending on production method; salmon byproducts typically contain 69-89% protein (3–5). FPH has been explored as a source of bioactive peptides with antioxidant, anti-hypertensive and anti-inflammatory effects (6, 7). However, according to statistics of Norwegian use, FPH is mainly used in pet food and feed formulations for farmed fish (1) rather than in human nutrition. The main barriers to consumer acceptance of FPH as food appears to be its fishy odors and flavors (1, 2, 8).

Consequently, there is a demand for novel strategies which enable the use of these nutritious fish byproduct-derived proteins in the higher-value food market. Several compounds are known to contribute to fish smell, of which trimethylamine (TMA) is a key component. TMA is a volatile, biogenic amine which is formed postmortem from trimethylamine-N-oxide (TMAO). TMAO is a naturally occurring metabolite in fish from cold and deep-sea environments and is, importantly, odorless. TMAO is believed to function as an osmolyte (9) and as a piezolyte, counteracting pressure-mediated inhibition of protein function (10, 11).

After slaughtering, TMAO in fish is converted to TMA by TMAO-reducing bacteria, contributing to its fish-like odor (12–14). The FPH industry has identified TMA as a key target for improving the sensory properties of FPHs. To remove it, some industrial actors currently use nanofiltration of FPH, but this untargeted method has the drawback of causing significant protein loss and altering the nutritional composition (15). An interesting alternative to filtration is to use an enzyme to specifically target TMA. Previously, we have shown that the bacterial trimethylamine monooxygenase mFMO can be used to remove most of the TMA from salmon FPH in a targeted approach (16, 17). The mFMO enzyme is a flavin-containing monooxygenase (FMO) (18) isolated from the marine gammaproteobacterium *Methylophaga aminisulfidivorans* (19). It belongs to a family of closely related bacterial FMOs which catalyze the oxidation of TMA (16, 20, 21). These bacterial FMOs oxidize TMA using molecular oxygen and NADPH as a cofactor, leaving TMAO and the oxidized cofactor NADP^+^ as products (Figure 1, top) (19, 21). To enhance compatibility with industrial processing conditions, a thermostable mFMO variant, termed mFMO_20, was generated through structure-based engineering and shown to perform well at up to 65°C (17). The oxidation reaction catalyzed by mFMO uses one molecule of NADPH per molecule of TMA. As NADPH is very expensive, mFMO-assisted removal of TMA in fish protein hydrolysates will not be industrially viable unless cofactor consumption is managed in a cost-effective manner.

**Figure 1:**
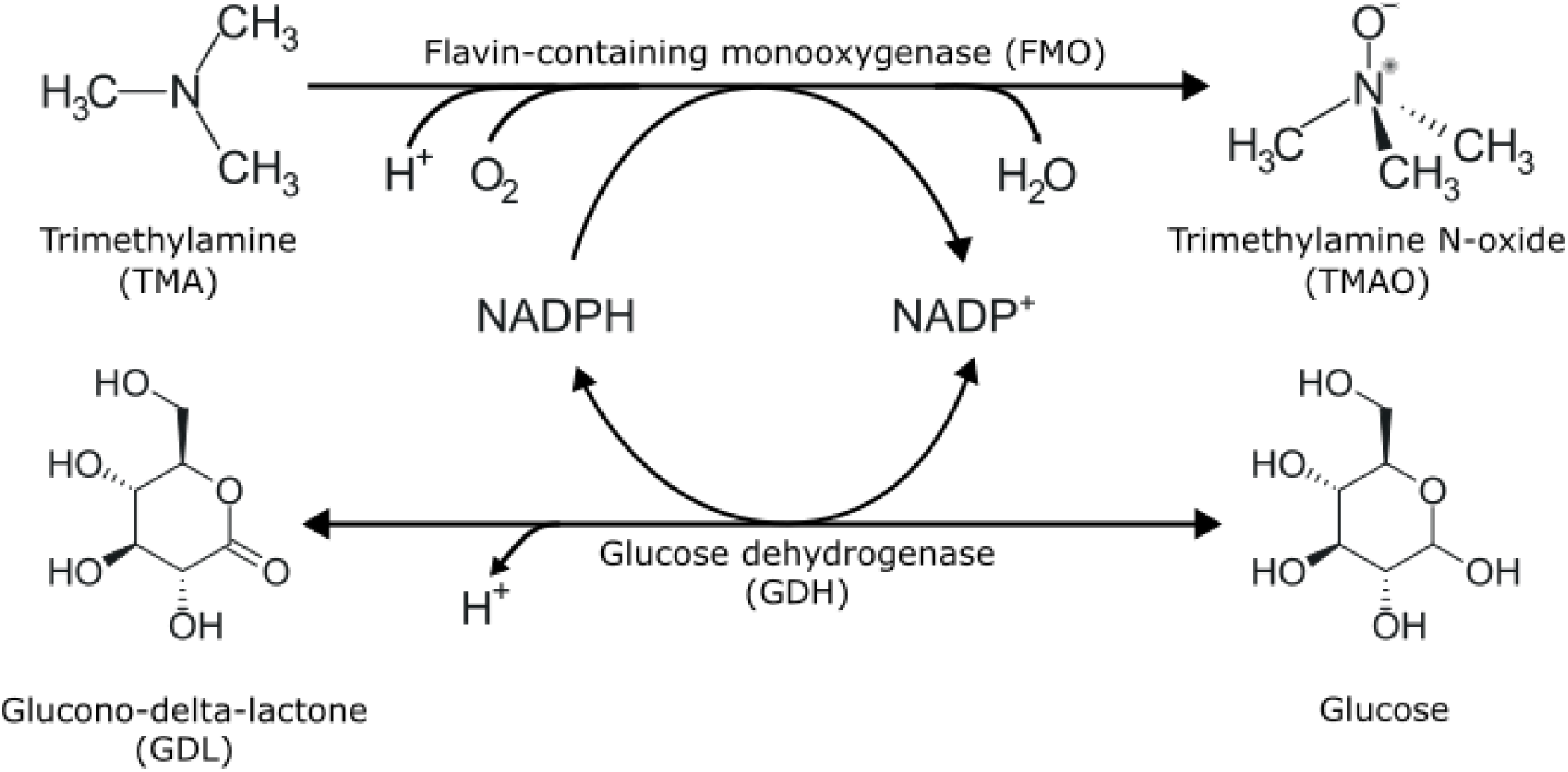
TMA oxidation and glucose dehydrogenation by FMO and GDH. Scheme of the TMA-to-TMAO and glucose-to-GDL reaction cycle. NADPH and NADP^+^, reduced and oxidized nicotinamide adenine dinucleotide phosphate, respectively.

Two strategies have been explored to reduce the cost of cofactors in reactions that depend on them: enzyme-cofactor engineering and cofactor recycling. Enzyme engineering can be used to improve the affinity for a cheaper cofactor with the same redox function, such as nicotinamide adenine dinucleotide (NADH) (22) or nicotinamide mononucleotide (23).

However, mutation may compromise other aspects of enzyme functionality, such as reaction rate and stability, requiring substantial screening to achieve a useful enzyme reaction rate (24). To our knowledge, successful attempts to change the cofactor specificity of FMOs from NADPH to NADH have not been reported. Furthermore, although NADH is cheaper than NADPH it remains prohibitively expensive for an industrial process (25). Cofactor recycling is thus an attractive strategy for cofactor management and has been successfully implemented in several systems (25). It involves regenerating the redox cofactor, enabling its reuse in multiple reaction cycles. This can be achieved through direct chemical reductive regeneration (26), homogenous (27, 28) and heterogenous (29) regeneration using hydrogen and organometallic catalysts, photocatalytic regeneration (30), and electrochemical regeneration (31). Alternatively, enzymatic regeneration (32–34) uses a secondary enzyme reaction, which reduces the cofactor while oxidizing a sacrificial substrate, to maintain cofactor availability and drive the main reaction. It has several advantages: it can be highly specific and, depending on the choice of sacrificial substrate, toxic components and catalysts can be avoided. (25). For food production involving enzymatic TMA-removal, the sacrificial substrate and products must food safe, and the regeneration enzyme must be compatible with both the processing conditions and the buffer requirements of the TMA oxidizing enzyme. Moreover, glucose is an inexpensive substrate, and this recycling strategy allows the use of the cheaper cofactor NADP^+^ instead of NADPH, significantly reducing costs associated with cofactor supplementation.

Various enzymes have been used to regenerate NADPH, including alcohol dehydrogenase, phosphite dehydrogenase, glucose dehydrogenase (GDH) and glucose-6-phosphate dehydrogenase (35). Phosphite dehydrogenase has been used in fusion constructs to enable sustained catalysis mediated by mFMO (36), and a Baeyer–Villiger monooxygenase (37). Like FMOs, the latter enzyme belongs to the class B flavoprotein monooxygenases (38). While GDH has not yet been reported for use with FMOs, it is an attractive cofactor-regenerating enzyme due to its widespread application, its simplicity, and the low cost of its substrate, glucose. Of note, glucose dehydrogenase B (GdhB) has been used to regenerate NAPDH during enzymatic Tyrian purple production by an FMO (39). For the application area in this study, it is important to note that both the glucose substrate and the product, glucono-1,5-lactone (also known as glucono-delta-lactone, or GDL), are recognized as safe and widely accepted food ingredients (Figure 1, bottom) (40).

In this study, we couple the activity of the thermostable mFMO_20 to the activity of the glucose dehydrogenase GdhB from *Priestia megaterium* (previously known as *Bacillus megaterium*) (41, 42). We demonstrate that this enzyme cascade, in the presence of excess glucose and the oxidized cofactor NADP^+^, effectively depletes TMA in salmon protein hydrolysate while recycling NADPH and producing GDL. Our approach demonstrates the utility of cofactor recycling for cost reduction in TMA remediation in an industrially relevant context.

## Results and discussion

### Establishing a cofactor recycling system for a TMA monooxygenase

When setting up a cofactor recycling system for enzymatic removal of TMA in FPH, we chose the GdhB enzyme and glucose as sacrificial substrate, and to use NADP^+^ as the added cofactor. To guide enzyme dosage and cofactor concentration in the dual enzyme system, we characterized the kinetics of GdhB for glucose. The K_m_ of GdhB for glucose was 68.7 mM (Table S1), a factor of 8 x 10^5^ higher than that of mFMO_20 for TMA (17). The V_max_ was about half that of mFMO_20. To compensate for the lower efficiency of GdhB, we set up the recycling system using a glucose concentration of 50 mM and a 10:1 ratio of GdhB to mFMO_20. Since NADH is more stable and less costly than NADPH, activities of a mFMO_20 and GdhB were assessed using both NADPH and NADH as electron donors. GdhB can catalyze oxidation of glucose by using both cofactors and can as well catalyze the reverse reaction using NAD(P)H and GDL (Supplemental Figure S1A). A sufficiently high glucose concentration is this required to drive the reaction towards GDL formation and NADP^+^ reduction. However, mFMO_20 does not accept NADH as a cofactor (Supplemental Figure S1B), necessitating use of NADP^+^.

To demonstrate that the mFMO_20/GdhB enzyme system catalyzed the expected reactions, we conducted an enzyme assay using the two enzymes and NADP^+^ as a cofactor. Two key features of the experimental design support this demonstration. First, we used NADP^+^ as a cofactor, rather than NADPH, ensuring that any TMAO production depended on GdhB activity. Second, TMA was added in a 5:1 molar excess relative to the cofactor, so that the formation of TMAO in quantities exceeding the initial cofactor concentration would provide direct evidence of cofactor recycling. The substrates and products were quantified using liquid chromatography/mass spectrometry (LC/MS) (Figure 2, Supplemental Table S3-S4). TMAO and GDL were formed when both enzymes, substrates and a cofactor were present. As expected, TMAO was not formed when any of the components were removed. When 100 μM NADPH was used as cofactor in the absence of GdhB, only 58 µM TMAO was formed, and as expected, no GDL was produced. Cofactor recycling was demonstrated by the formation of 191 µM TMAO a reaction with only 100 µM NADP^+^. This confirms that cofactor recycling enabled the oxidation of TMA in molar amounts exceeding the initial cofactor concentration.

**Figure 2:**
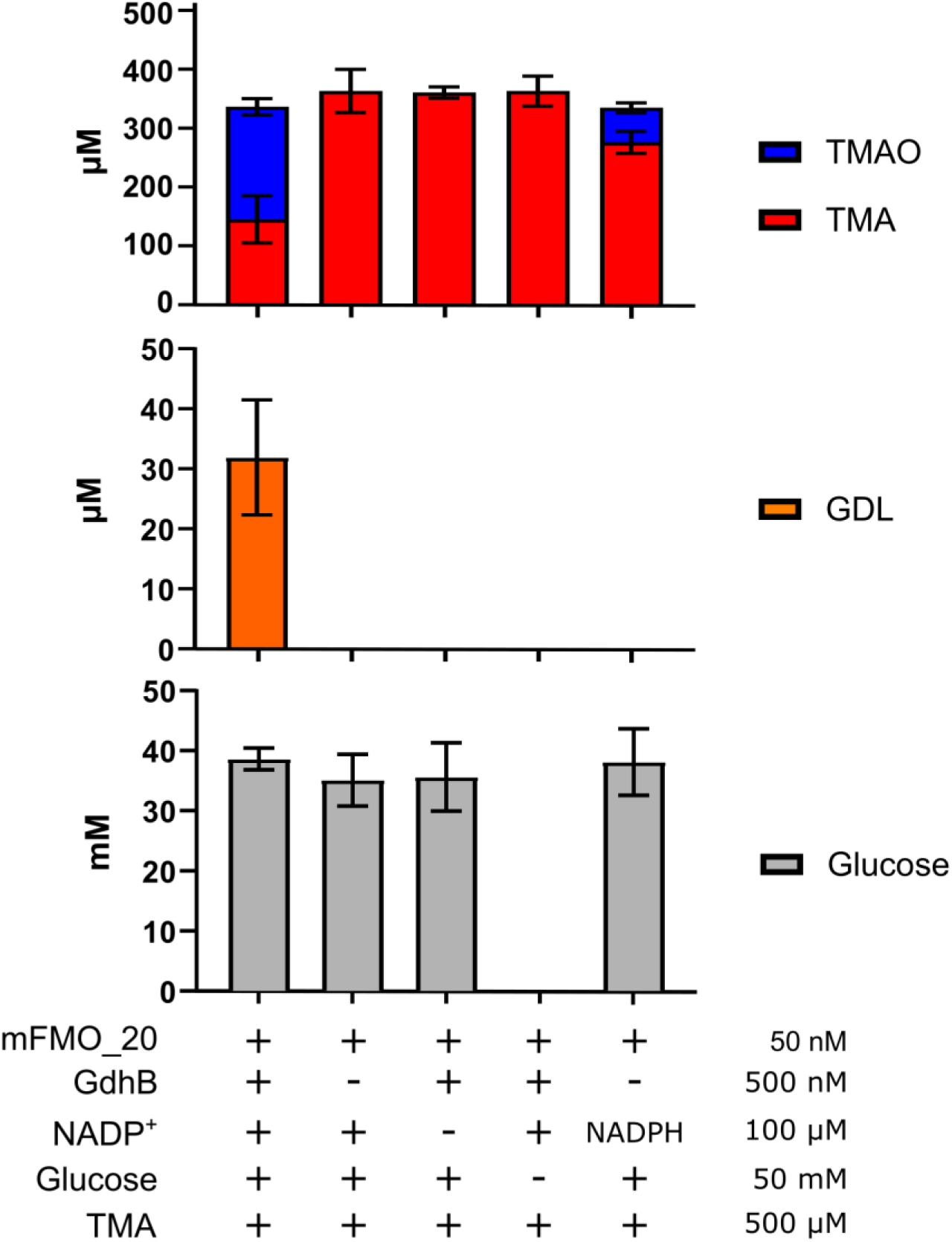
TMA oxidation driven by cofactor recycling. Enzyme assays (n=3) with the indicated components (+, present; -, absent): mFMO_20, GdhB, NADP^+^ (or NADPH), glucose and/or TMA at the given concentrations, were incubated for 1 hour at 25°C and analyzed by LC/MS for the presence of TMA, TMAO, glucose or GDL, which were quantified against a standard curve.

The GDL concentrations were lower than expected, as we anticipated GDL and TMAO to increase in concert. This discrepancy may be due to spontaneous hydrolysis of GDL to gluconic acid, a process reported in the literature (40). Although this possibility was considered during LC/MS method development, no ions corresponding to gluconic acid were detected. A disadvantage of using GdhB for cofactor recycling in this system is its relatively low catalytic activity compared to mFMO_20, necessitating a relatively high concentration of glucose to drive the recycling reaction. We used 50 mM glucose and 100 μM NAD^+^, which is consistent with the 100 mM glucose and NAD^+^ concentrations of 10-500 mM used in previous studies with this enzyme (41, 43, 44). The final glucose concentration in the protein hydrolysate obtained in this study was relatively high at 1.75% w/w, assuming a 10% dry weight content (see next section). However, this level is comparable to the 1% xylose concentration used with heat to achieve browning and caramelized flavor for odor masking of salmon protein hydrolysate (45). In comparison,). To achieve better TMA remediation with lower levels of glucose in the final product, it could be useful to improve the activity and substrate affinity of GdhB through enzyme engineering (46) or through immobilization (47, 48).

### The dual enzyme system reduces TMA levels in salmon protein hydrolysates

To demonstrate that the mFMO_20/GdhB enzyme cascade can deplete TMA in an industrially relevant protein hydrolysate, a lab-scale protease-driven hydrolysis of salmon heads and frames (byproducts) was performed (Figure 3).

**Figure 3:**
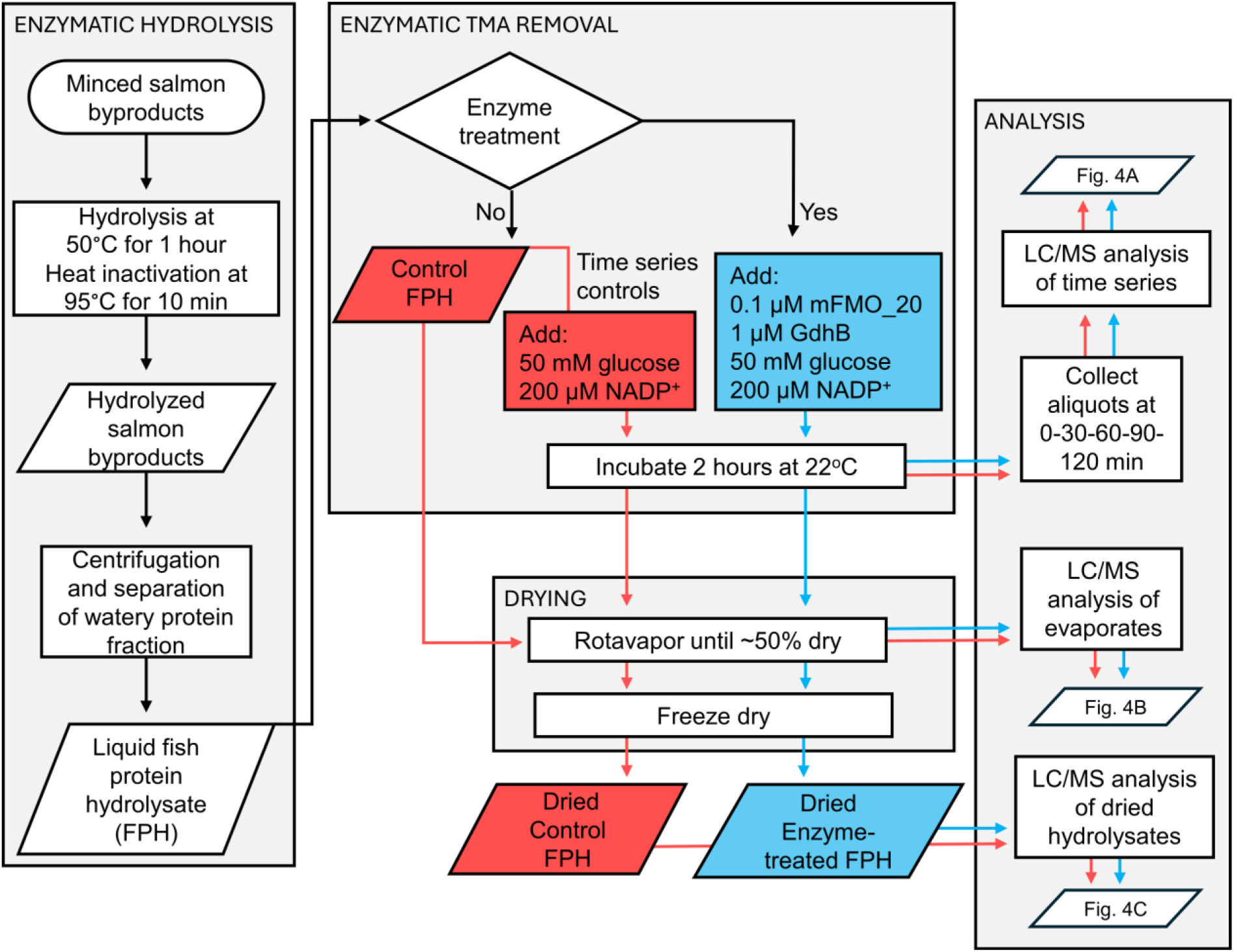
Production and evaluation of enzyme-treated FPH at lab scale. The rectangles indicate process steps and analyses, while diamonds show branching points, and the parallelograms contain intermediate and final products, and indicates which figure contains the associated result. The flow of control FPH fractions are shown in red and enzyme-treated FPH fractions in blue. FPH: fish protein hydrolysate (salmon); LC/MS: liquid chromatography/mass spectrometry.

Whole, fresh salmon were filleted, and the heads and frames were minced in a meat grinder to produce the byproduct feedstock. This mince was mixed with 50% water (w/w) and 0.5% Alcalase 2.4L (v/w biomass), a commercial subtilisin endoprotease with broad specificity. After hydrolysis and heat inactivation of the protease, the sample was centrifuged to isolate the water-soluble fraction containing hydrolyzed peptides, hereafter referred to as FPH. To protect the added mFMO and GdhB enzymes from proteolytic cleavage and lipid interference, they were introduced only after hydrolysis and centrifugation, along with glucose and NADP^+^ (Figure 3). We obtained a total of 4860 ml liquid hydrolysate which served as the substrate for enzymatic TMA removal treatment (Figure 3, top middle box). The dry matter content was estimated to be 9.3%.LC/MS analysis of FPH samples collected during the enzymatic TMA removal process showed that treatment with mFMO_20 and GdhB depleted TMA in a time-dependent manner, reducing it to less than 1% of the initial intensity (Figure 4A, Supplemental Table S5). The TMA intensity was unchanged in controls where enzymes, glucose, and NADP^+^ were absent. The TMAO intensity was stable throughout the time course, likely reflecting the high TMAO content in the freshly prepared FPH. The glucose concentration also remained stable, as it was added in great excess. GDL intensity increased over time in parallel with the reduction of TMA, confirming successful cofactor regeneration. The final dried control and enzyme-treated FPHs were prepared by evaporation until partially dried, followed by freeze drying until completely dry. Analysis of the evaporate (Figure 4B, Supplemental Table S6) revealed approximately 50 µM TMA remaining in the control FPH, while no TMA was detected in the enzyme-treated evaporate. This demonstrates that although drying may assist in TMA removal it is not sufficient to fully deplete it. Quantification of TMA in the dried FPHs showed that 58 ppm remained in the mFMO_20/GdhB-treated FPH, whereas 208 ppm was retained in the control (Figure 4C). Hence, the enzyme treatment oxidized approximately 75% of the TMA content. A previous study reported that the application of nano- and diafiltration reduced TMA from 700 ppm to 100 ppm in cod FPH and from 400 ppm to 100 ppm in salmon FPH (15). Reaching a level of 100 ppm was associated with improved TMA taste intensity. We did not observe an increase in TMAO concomitant with the decrease in TMA in the enzyme-treated FPH. The 150 ppm TMAO produced may not make enough of a difference to be detected between samples, given that the TMAO concentration was measured between 1344 and 1615 ppm, and the variation between replicate injections was between 4.9 and 12.2% (Supplemental Table S6). Neither did we detect GDL in the enzyme-treated FPH. The lower limit of quantitation of GDL was 4.2 pmol, while the reduction in TMA was measured at 7.62 pmol (150 ppm). Given that the TMA conversion exceeded the GDL quantitation limit, we would have expected to detect GDL. One possible explanation for this apparent discrepancy is hydrolysis of GDL into gluconic acid, which would prevent GDL accumulation and detection.

**Figure 4:**
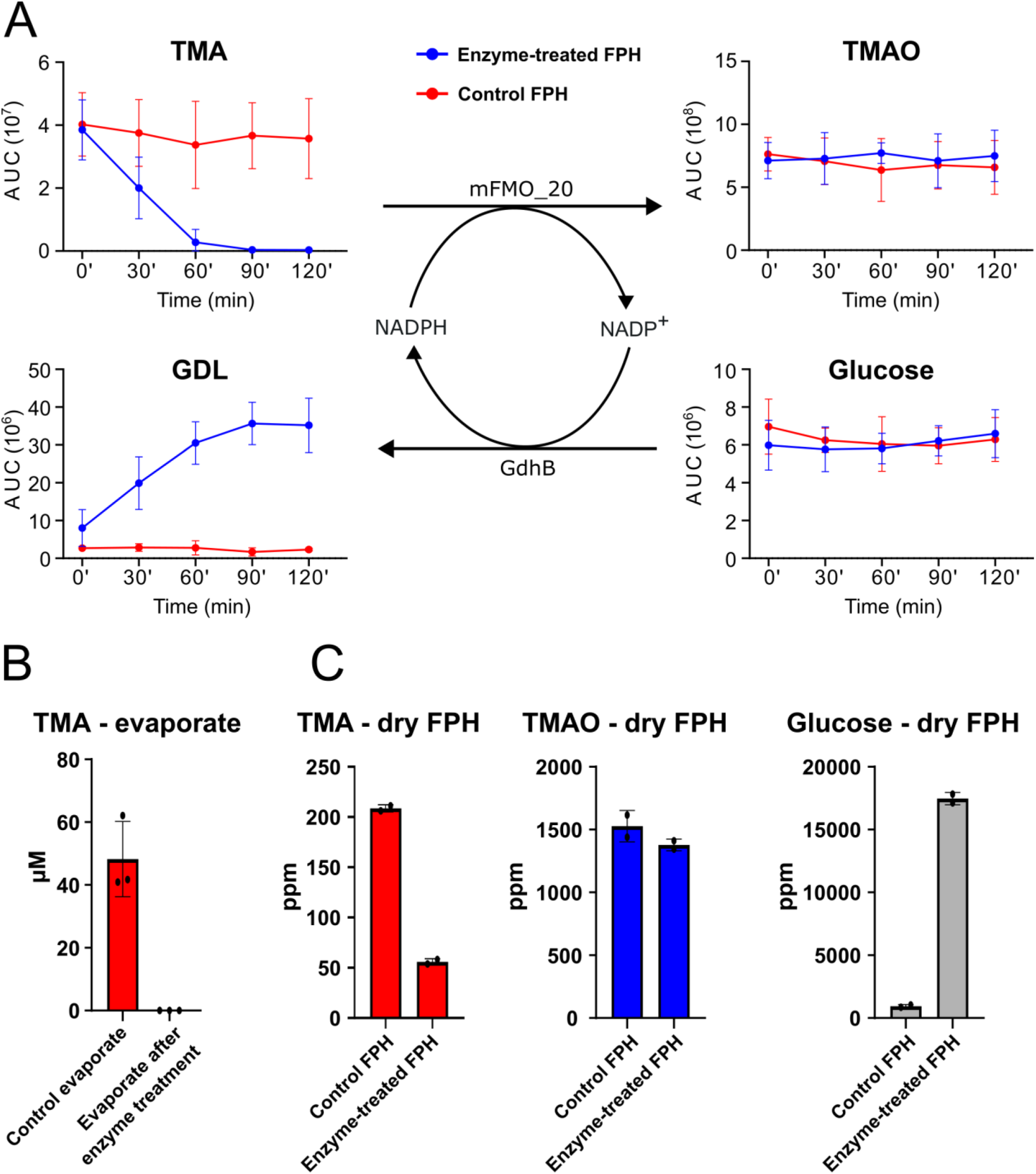
Cofactor recycling-driven TMA oxidation in salmon protein hydrolysate. A) Time course analysis of the enzymatic TMA removal process. TMA, TMAO, glucose and GDL were quantified at each time point by LC/MS. Blue: enzyme-treated FPH (presence of mFMO_20/GdhB); red, control FPH (absence of mFMO/GdhB). n=6 for each time point (0, 30, 60, 90, 120 min) for control FPH and n=4 for enzyme-treated FPH. Error bars show standard deviation between replicates. AUC: area under the curve. B) TMA in the enzyme-treated and control FPH evaporates measured by LC/MS (n=3). Error bars show standard deviation between evaporate batches. C) TMA, TMAO, glucose and GDL quantified by LC/MS in dried enzyme-treated FPH and control FPH; n=2 injections of the same sample. Error bars show standard deviation.

### Reduction of TMA levels in FPH assessed by sensory panels

To assess whether the reduction in TMA measured by LC-MS corresponded to a decrease in smell intensity, a trained sensory panel evaluated multiple smell attributes of the dried control and enzyme-treated FPHs. The evaluation included TMA and overall smell intensity, rated on a 9-point scale (Figure 5 and Table S7). The assessors detected significantly lower intensities in nine of the ten attributes for the enzyme-treated FPH. Notably, the TMA smell was perceived to be significantly lower by 2.5 units, which is in line with the observed 75% reduction in TMA levels (Figure 4). The sensory panel is trained to detect TMA as distinct from other aromas such as rancid or seaweed smells. Hence, the reduction in other sensory attributes is probably not caused by the reduction in TMA levels and may instead reflect enzyme activity on other compounds or effects from glucose.

**Figure 5:**
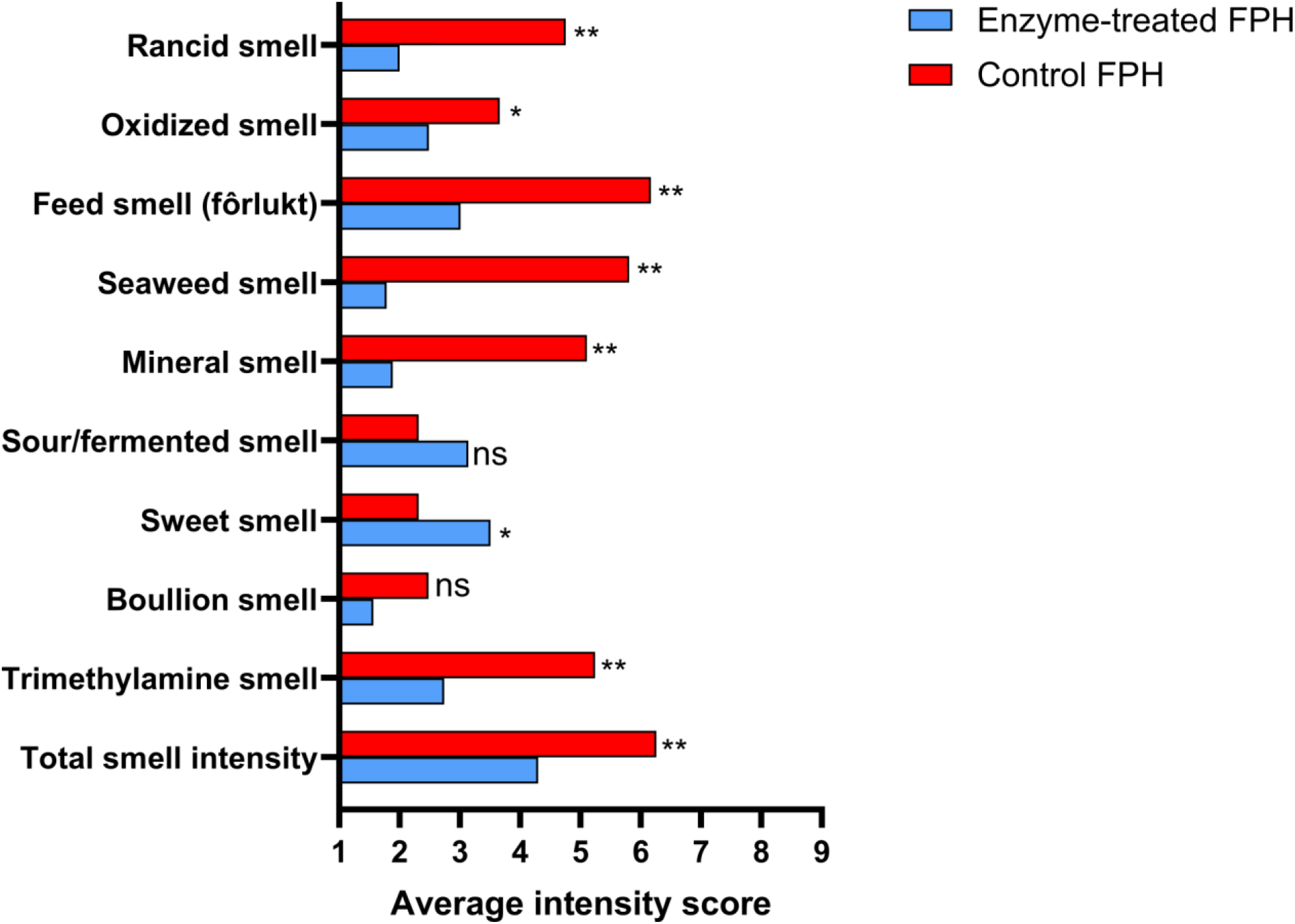
Assessment of fish protein hydrolysates by a trained sensory panel. Each sensory panel member (n=8) scored the intensity of each dried hydrolysates twice on ten smell characteristics in a blinded and randomized order. Enzyme-treated FPH and control FPH indicates presence and absence of mFMO_20/GdhB treatment, respectively. Statistical significance was tested using ANOVA and Tukey’s test for multiple comparisons. **: p<0.01, *: p<0.05, ns: p>0.05. See Table S3 for description of each attribute.

Notably, the enzyme-treated FPH scored significantly higher on sweet smell, likely due to the added glucose absent in the control. Both native mFMO, which served as the basis for mFMO_20 used in this study, and FMO from *Methylocella silvestris* have been reported to oxidize compounds beyond TMA, including dimethylamine, methimazole, indole and dimethyl sulfide (19, 21). Although off-target oxidations by mFMO_20 have not yet been investigated, the oxidation of unknown metabolites may have influenced the olfactory profile of the FPH.

The ultimate aim of the TMA remediation process is to reduce TMA levels in FPH sufficiently to achieve consumer acceptance. To evaluate whether this reduction translates into improved consumer acceptance, 70 participants aged 24-59, of whom 61% were women, were recruited from the Province of Barcelona, Spain, for preference and acceptance testing of the enzyme-treated, dried FPH (Figure 6). Participants evaluated acceptance, using a 7-point hedonic scale ranging from “dislike very much” (1) to “like very much”, and rated smell intensity on a 7-point scale from lowest (1) to highest (7). The intensity attributes assessed were fishy smell, ammonia-like smell, sulfur-like smell, freshness, and overall smell intensity. Both the control and enzyme-treated FPHs showed similar acceptability and attribute intensities. The enzyme-treated FPH had an average acceptability score of 3.2, while the control scored 3.5, with a median of 3 for both (Figure 6A). Average scores for fish smell attributes were not markedly different between the samples (Figure 6B). Respondents were also asked to rank the samples by preference. Of the 69 respondents who indicated a preference (one did not), 42 (60.9%) preferred the control FPH, while 27 (39.1%) favored the enzyme-treated FPH. We conclude that the consumer panel did not perceive a difference between the enzyme-treated and control FPHs. The discrepancy between the two panels suggests that trained assessors are more sensitive to subtle changes in specific odor attributes in FPH due to their calibration and experience. In contrast, consumer perceptions may be more variable and influenced by overall impression or other sensory cues. This highlights the importance of using both types of panels to gain a comprehensive understanding of sensory changes.

**Figure 6:**
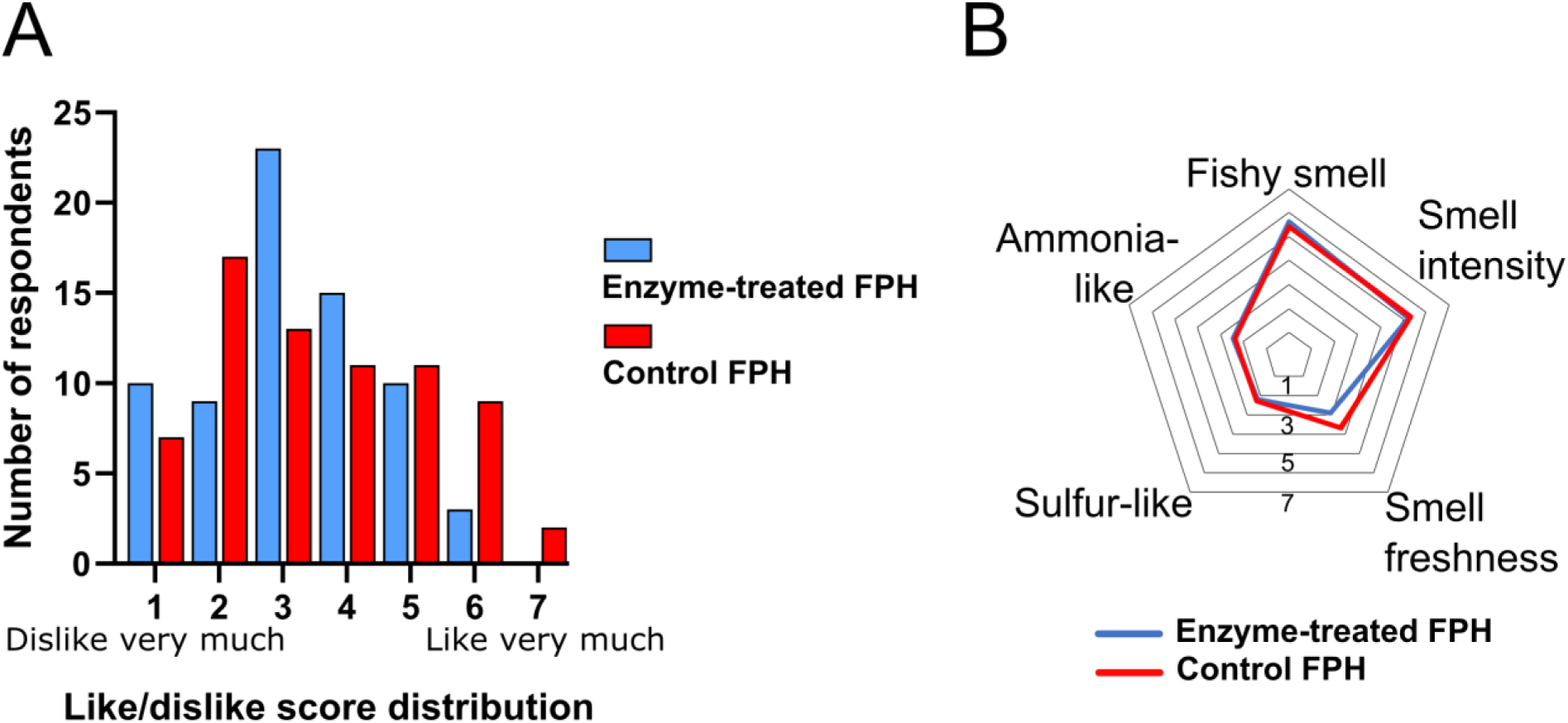
Sensory assessment of enzyme-treated FPH by a consumer panel. A) Consumer acceptability test with distribution of smell perception scores. Participants (n=70 participants) rated the smell of each hydrolysate on a 7-point scale, where 1 corresponds to “dislike very much” and 7 to “like very much”. The description of the scale is given in Supplemental Table S8. B) Average intensity scores were obtained from consumer panelists (n=70) who rated the smell of each hydrolysate based five attributes: fishy smell, ammonia-like smell, sulfur-like smell, smell freshness, and smell intensity. Ratings were given on a 7-point scale, where 7 indicated the highest intensity. Descriptions of the attributes are given in Supplemental Table S9.

TMA content of seafood and its raw materials varies with fish species and freshness. TMA is reported to vary between 0.3 g/kg wet weight (49) and 0.7 g/kg dry weight (15) in cod FPH, and between 0.4 g/kg (15) and 1.1 g/kg dry weight (17) in salmon FPH. Converting 1 gram of TMA in FPH would require over 12 grams of the reduced cofactor, NADPH, and would thus be economically prohibitive.

The final TMA concentration of the mFMO_20/GdhB-treated FPH (58 ppm) remains five orders of magnitude higher than the reported human detection threshold for TMA (0.00021 ppm v/v) (50). This suggests that the TMA smell would be detectable even after enzyme treatment. While removing TMA was expected to improve organoleptic properties by reducing TMA-related sensory attributes, the residual TMA level indicates that further optimization is needed. Additional experiments are needed to investigate if cofactor recycling can sustain the mFMO catalysis for longer and fully deplete TMA. A GdhB-driven NADPH regeneration system sustained the production of (R)-4-chloro-3-hydroxybutanoate ethyl for five hours (34). However, a continuous reaction may depend on factors such as NADP^+^ dimerization or isomerization causing cofactor loss (25), pH changes that affect enzyme activities, or product inhibition.

Although TMA is widely recognized as a key contributor to the characteristic odor of fish and FPH (8), it is likely not the sole determinant of what is perceived as fishy smell. The overall sensory perception is likely influenced by a complex mixture of volatile organic compounds, including aldehydes, ketones, other amines, and sulfur-containing molecules, and concentrations and volatility also play a role (51, 52). A combination of targeted and untargeted approaches, for example filtration combined with enzyme treatment, may be needed to obtain FPH products that are acceptable to end users. It remains uncertain whether reducing TMA concentrations to imperceptible levels alone will sufficiently diminish the overall fishy smell.

### Conclusion and future opportunities

We have shown that an enzyme cascade combining mFMO and GdhB serves as a targeted approach to reduce TMA content in FPH, while lowering costs through efficient cofactor recycling. Using a combination of quantitative instrumental and sensory approaches, our results demonstrate both a decline in TMA levels and a corresponding reduction in TMA-derived sensory attributes. These results suggest that, with further optimization of reaction time and conditions, enzymatic treatment has the potential to achieve greater depletion of TMA than untargeted approaches. To develop palatable FPH products, further research is needed to unravel the role and complexity of odor-contributing components, as the mechanisms underlying fishy smell formation are not yet fully understood. Exploring approaches to produce FPH that meet end-user acceptance offer an avenue for future investigation.

## Methods

### Fermentation and purification of enzymes

mFMO_20 and GdhB enzymes were expressed in *E. coli* from a vector encoding a C-terminal His-tag. GdhB additionally carries an N-terminal SpyTag (53). GdhB was expressed in 10 L 2YT medium (16 g/L tryptone, 10 g/L yeast extract, and 5 g/L NaCl) mFMO_20 was expressed in a in 8 L medium with the following composition (in g/L unless otherwise specified): Na_2_HPO_4_ • 7 H_2_O, 12.8; KH_2_PO_4_, 3.0; NaCl, 0.5; NH_4_Cl, 1.0; MgSO_4_ • 7H_2_O, 0.5; CaCl_2_ • 2 H_2_O, 0.015; Glucose, 0.5; Ampicillin,0.1; Casamino acids, 10; Glycerol, 4.25; L-leucine, 0.00005; trace metal solution, 1 ml. Composition of trace metal solutions (in g/L) FeCl_2_ • 4 H_2_O,1.5; ZnCl_2_, 0.07; CoCl_2_ • 6 H_2_O, 0.19; MnCl_2_ • 4 H_2_O, 0.1; CuCl_2_ • 2 H_2_O, 0.02; NiCl_2_ • 6 H_2_O, 0.024; Na_2_MoO_4_ • 2 H_2_O, 0.036; H_3_BO_3_, 0.006; 25% HCl, 10 ml. Both enzymes were expressed in a 15 L turbine-stirred bioreactor (Chemap, Switzerland). Cultivation of both enzymes was started at 37 ± 0.1°C. A pH meter (Mettler Toledo InPro® 3030/120) was used to maintain the pH at 7.0 ± 0.1 with 1 M NaOH and 1 M H_2_SO_4_. The fermenter was operated at an overpressure of 0.2 bar and with constant supply of sterile air (2.5 L/min) through the bottom sparger. Dissolved oxygen was maintained at minimum of 30% of air saturation by regulating the stirrer speed during cultivation. Foaming was controlled by dropwise addition of an anti-foaming agent (A240, Sigma). Percentages of O_2_ and CO_2_ in the off gas were measured continuously using a mass spectrometer (Prima Pro Process Mass Spectrometer, ThermoFisher) and the experimental data was acquired from LabVIEW 6 (National Instruments, USA). The batch phase started by inoculating the media with 500 ml of inoculum. When the optical density at 600 nm reached between 0.5 – 0.8, the cultivation temperature was reduced to 18 – 20°C and the cultures were induced with 20% L-arabinose (final concentration in the broth 1 g/L) to initiate the enzyme production. When the fermenter off–gas CO2 concentration dropped, the batch phase was over. After completion of the batch experiment, the fermentation broth was harvested. The cell biomass was separated from the broth by centrifugation at 4500 rpm for 20 min at 4°C. Cell paste was stored at - 20°C. mFMO_20 was purified essentially as described (17) in 50 mM tris-HCl, pH 8.0 and 500 mM NaCl, while the buffer system for GdhB was 20 mM Na_2_HPO_4_-NaOH, pH 6.5 and 500 mM NaCl. Equilibration and washing buffers were supplemented with 10 mM and 50 mM imidazole, respectively. The elution buffer for mFMO_20 and GdhB contained 500 mM and 300 mM imidazole, respectively. The desalting buffers consisted of 500 mM NaCl and 10% glycerol. Cells were lysed in lysis buffer (equilibration buffer supplemented with 25 µg/ml lysozyme and 1 mM PMSF) by three freeze/thaw cycles followed by sonication on ice (1/8” probe, 70% amplitude, 10 seconds on and 20 seconds off, for a total of 2 minutes sonication time).

mMFO_20 was purified using an Äkta Pure chromatography system (GE Healthcare) with a 5 ml His-Trap column (GE Healthcare). The lysate was cleared for 30 minutes at 17,000 x *g* and loaded on the column using the sample pump. After washing with 20 column volumes (CVs) of wash buffer, the His-tagged protein was eluted by a stepped gradient of 5 CVs per step, at 35%, 50%, 65% and 100% B. The eluate was collected in 2 ml fractions. Fractions which appeared yellow, likely due to bound FAD (54), were combined and buffer was exchanged to desalting buffer by sequential centrifugation using a 30 MWCO filter. Enzymes were aliquoted after concentrating to 6.9 mg/ml, and stored at -80°C.

GdhB was purified using benchtop columns with 1.5 ml Ni-NTA resin (Qiagen). Cells were lysed and lysate was cleared in the same manner as for mFMO_20. The lysate was loaded onto equilibrated benchtop columns by gravity flow. Columns were washed with 20 CVs wash buffer and eluted in 2.5 ml elution buffer. After elution, GdhB was desalted using PD-10 columns (Cytiva) equilibrated with desalting buffer and eluted with 3.5 ml desalting buffer.

The concentration at this point was between 1-3 mg/ml. GdhB was then aliquoted and kept at-80°C until use. Purity was analysed by visual inspection by sodium dodecyl sulfate polyacrylamide gel electrophoresis. Concentration was measured using the 660 nm protein assay kit (Pierce), quantified against a standard curve of BSA.

### Enzyme assays and enzyme kinetics

Enzyme activity of mFMO_20 and GdhB was measured by spectrophotometrically monitoring the oxidation of NADPH or NADH, using the change in absorbance at 340 nm, and assuming a molar extinction coefficient of 6220 M^-1^ cm^-1^. Enzymes were diluted to 1-10 µM and used at a final concentration of 50-1000 nM, mixed with cofactor (NAD(P)H or NAD(P)^+^), and 100 µM TMA or 50 mM glucose or GDL. Activity was calculated as described (17). Briefly, all components except substrate were mixed and diluted to 1 ml in a polystyrene cuvette, using 50 mM tris-HCl, pH 8.0 and 50 mM NaCl as the reaction buffer, and the reaction was started by adding the relevant substrate (TMA, glucose or GDL).

Absorbance at 340 nm was monitored in a Cary UV-VIS photospectrometer over 1 minute for mFMO_20 and 2 minutes for GdhB. The absorbance change per minute (positive for the NADP^+^ reduction reaction and negative for the NADPH oxidation reaction) was used to calculate the cofactor concentration change, assuming a molar extinction coefficient of NADPH of 6200 M^-1^ cm^-1^. Activity is expressed as µM product formed min^-1^ µM enzyme^-1^, or min^-1^. Kinetic characterization of GdhB with constant NADP^+^ (100 µM) was performed with 0.1-1000 mM glucose, in duplicate measurements. The substrate concentration [mM] versus velocity [min^-1^] was plotted and the Michaelis-Menten parameters of K_m_ and V_max_ were calculated in GraphPad Prism.

### Cofactor regeneration enzyme assays

mFMO_20 (final concentration: 50 nM), GdhB (500 nM), cofactor (NADP^+^ or NADPH, 100 µM), glucose (50 mM) and TMA (500 µM) were diluted in 50 mM NaCl and 50 mM tris-HCl, pH 8.0. As reaction controls, either GdhB, NADP^+^ or glucose were omitted. The total reaction volume was 50 µl in a 1.5 ml microcentrifuge tube, and each condition was repeated 3 times. The reaction was started by addition of substrate – glucose, or in the case of the condition with NADPH, by addition of TMA. Reactions were briefly vortexed and centrifuged, and reactions were incubated at 25°C with light agitation for 1 hour. The reactions were stopped by addition of 450 µl 100% LC/MS-grade acetonitrile, vortexed, and stored at -20°C until analysis by LC/MS.

### FPH production by enzymatic hydrolysis

Two fresh, gutted salmon (10.186 kg) were purchased (Meny, Bergen Storsenter), and filleted. The remaining byproducts consisted of 2.430 kg of salmon heads, backbones, fins, tails, and skin (giving a byproduct fraction of 23.9%). Byproducts were minced and frozen at -20°C. Enzyme-assisted hydrolysis was performed in a customized bioreactor (H.E.L. automate, UK) by mixing 250 g thawed, minced byproduct with 250 ml water per reaction vessel, and heating to 50°C before adding 0.5% (w/w) Alcalase 2.4L (Sigma-Aldrich) and allowing hydrolysis to proceed for 1 hour at 50°C. The enzymes were heat inactivated at 90°C for 10 minutes. Insoluble material was removed by centrifugation at 5000 x *g* and 15000 x *g* for 20 minutes at 4°C. The water phase was separated from the oil phase using a separation funnel. The water phase was filtered through a 7 µm filter paper, then through 1.2 µm and 0.45 µm cellulose acetate filters. The filtered water phase was stored at -20°C.

### Enzyme treatment of FPH

For enzyme treatment, filtered FPH (in total 2.4 L, from 8 x 300 ml batches) was mixed with mFMO (final concentration: 100 nM), GdhB (final concentration: 1 µM), glucose (final concentration: 50 mM), and NADP^+^ (final concentration: 200 µM), and allowed to react for 2 hours at 25°C under agitation. Samples for LC/MS analysis (50 µl) were taken after 0, 30, 60, 90 and 120 minutes. In parallel with each batch, 30 ml of hydrolysate controls without enzymes, cofactor or glucose were incubated for timepoint controls. To dehydrate the hydrolysate, it was dried in a Rotavapor set to 60°C and 150-125 mbar of vacuum, until the volume was 14-32% of the starting volume. The hydrolysate was further dried to completion in a freezer dryer, and the batches were combined and ground to a powder using a mortar and pestle. The non-enzyme treated hydrolysate (2.05 L, dried in 7 batches of approximately 300 ml, were dehydrated in the same manner.

### Analytical evaluation of FPH and enzyme assays

TMA conversion and the glucose dehydrogenase reaction were monitored by LC/MS. To prepare samples for LC/MS analysis, 50 µl sample is mixed with 450 µl 100% acetonitrile and vortexed, then centrifuged for 20 minutes at 17,000 x *g* and 4°C. 75 µl of the supernatant is then transferred to an autosampler vial and kept at 4°C until analysis. The standards were prepared by serial dilution in 90% acetonitrile. The dried FPH was diluted to 1 mg/ml in 90% acetonitrile before clearing.

The mobile phase is 10 mM ammonium acetate, pH 8.0. Buffer A has 3% acetonitrile and buffer B has 90% acetonitrile. The column is an ACQUITY Premier BEH Amide VanGuard FIT HILIC column (Waters), with particle size 1.7 µm and internal dimensions of 2.1 x 100 mm, fitted with a 2.1 x 5 mm guard column. 3 µl sample or standard were loaded. The method takes 14 minutes and uses a 0.2 ml/min flowrate, starts at 85% B, going to 80% over 1.6 minutes, then to 40% at 10 minutes, holding at 40% until 12.7 minutes, while final conditioning at 80% B happens between 12.8 and 14 minutes. An Orbitrap Q-Exactive (Thermo Fisher) is connected online to a Dionex UltiMate 3000 UHPLC system (Thermo Fisher). The mass spectrometer is operated in the positive mode for full ion scan, with scheduled switching to negative mode when GDL elutes at approx. 2.2 minutes.

Quantification of the selected ions is performed in Excalibur Quant Browser (Thermo Fisher) by taking the area under the curves of the extracted ion chromatograms from samples and standards in defined RT windows (Table S2). The enzyme treatment of the hydrolysates was performed in batches, and these batches served as replicates for the statistical treatment of the quantified ions.

### Sensory assessment by trained panel

Dried enzyme treated and control FPH were assessed by a trained sensory panel at Nofima AS, using a Quantitive Descriptive Analysis (ISO 13299:2016) and according to a Generic Descriptive Analysis (55). The sensory panel consists of 8 judges, trained and satisfying the requirements of ISO 8586-1:2012. The rooms where the test was conducted were built according to ISO 8589:2007, and have individual judging booths, standardized lighting, and its own ventilation system. The judges were presented with the dried hydrolysates (either control or enzyme-treated), blinded as to which sample was which. Samples were served in a blocked and randomized fashion to each judge. 1 gram of sample in white cups with metal lids were served. They were evaluated on the smell parameters of total smell intensity, trimethylamine smell, boullion smell, sweet smell, sour/fermented smell, seaweed smell, feed smell, oxidized smell, and rancid smell. The parameters are described in Table S3. Each judge scored the intensity of the samples 1-9 for each parameter, with 1 being least intense and 9 being most intense. Panel averages were compared by ANOVA, and correction for multiple testing was done by Tukey’s test. Difference between control and enzyme-treated FPH was deemed significant if the corrected p-value was <0.05.

### Consumer recruitment for sensory testing

A total of 70 adult consumers were recruited from among the employees of Leitat Technological Center, located in the Province of Barcelona (Spain), to take part in a sensory evaluation focused on odor perception. Recruitment was carried out internally by sending an email invitation to staff member through the organization’s internal mailing system. The email included a brief description of the study’s objective, the nature of the sensory evaluation (limited to smelling two samples), and the criteria for participation. Eligible participants were adults (≥18 years) with no known olfactory impairments. Employees interested in participating were directed to an online registration form, which collected demographic information (age, gender, location), verified eligibility, and allowed individuals to indicate preferred time slots. Although no strict quotas were applied, efforts were made to ensure demographic diversity among the participants. Before the evaluation, all participants signed an informed consent form, in compliance with ethical research standards approved by the relevant institutional review board.

### Consumer panel evaluation

During the sensory session, each participant was presented with two blinded hydrolysate samples contained in odor-isolated vessels. Sample A (enzymatically treated) and sample B (no treatment). Both samples have been prepared under the same conditions, ensuring uniformity in quantity and presentation and have been given to the panelists in a room at a constant temperature and isolated from external odors to avoid interferences. Panelists evaluated specific odor-related parameters: fish smell, smell intensity, smell freshness, sulfur like smell and ammonia like smell, using intensity and acceptability scales of seven points. The parameters are described in Table S4, and the intensity descriptions are in Table S5. A preference test between samples was also performed (56). Consumers were asked to rank the samples in order of preference. To obtain the ranking of each sample, each rank position was multiplied by the number of consumers that had selected it, and the sum of the rankings of each sample was calculated. Low values in rank sum of samples indicated that the sample has mainly been ranked in the first order of preference.

## Acknowledgements

All authors received funding from the European Union’s Horizon 2020 research and innovation program under Grant Agreement 101000607 (OXIPRO). LC/MS analyses were performed at the Department of Bioscience, University of Bergen, and the authors thank Ersilia Bifulco (University of Bergen) for support with the LC/MS data collection. Sensory assessment by the trained panel was performed at Nofima AS.

## Supplementary figures and tables

**Figure S1:**
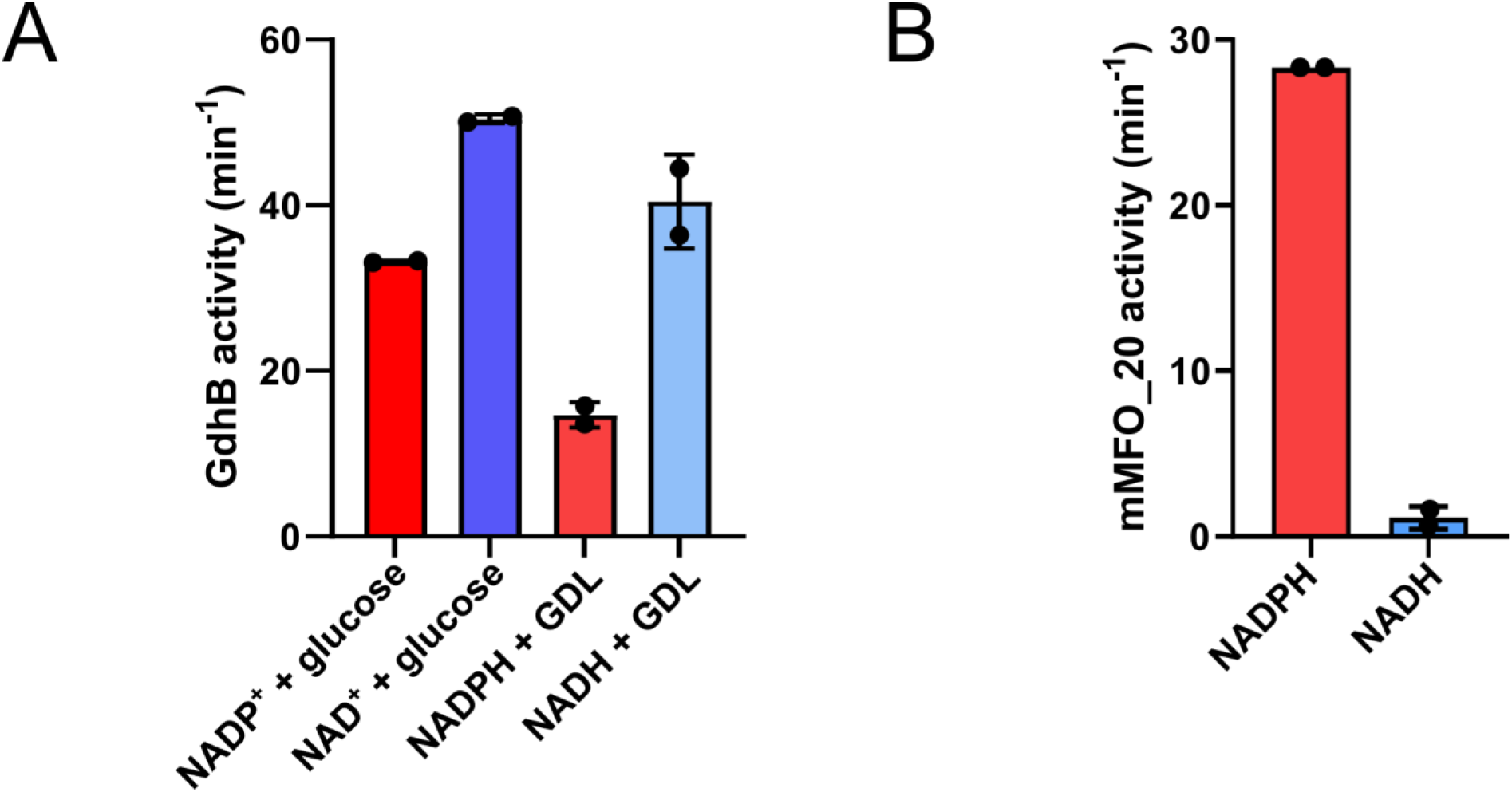
A) Activity of purified GdhB (500 nM) with glucose (50 mM) and the indicated oxidized cofactor (100 µM), and with the indicated reduced cofactor and GDL (50 mM), measured by absorbance change at 340 nm. B) Activity of purified mFMO_20 (50 nM) with 100 µM trimethylamine (TMA) and 100 µM reduced NADPH or nicotinamide adenine dinucleotide (NADH), measured by absorbance change at 340 nm.

**Table S1:** Kinetic parameters of GdhB using glucose as a substrate. NADP^+^ concentration. Parameters of mFMO_20 are included for reference.

| Enzyme | $V_{\max}$ (s<sup>-1</sup>) | $K_m$ (mM) | Specificity constant (s<sup>-1</sup> M<sup>-1</sup>) | Reference |
| --- | --- | --- | --- | --- |
| GdhB | 0.429 | 68.68 (glucose) | 6.25 | This study |
| mFMO_20 | 0.93 | 0.83 x 10<sup>-3</sup> (TMA) | 1.11 x 10<sup>6</sup> | Goris et al, 2023 |

**Table S2:** Metabolites and related analytical information quantified by LC/MS.

| Metabolite | Molecular formula | Mass (g/mol) | Ion (m/z, charge) | Retention time (minutes) |
| --- | --- | --- | --- | --- |
| TMA | C<sub>3</sub>H<sub>9</sub>N | 59.112 | 60.08160 (+H) | 3.82 |
| TMAO | C<sub>3</sub>H<sub>9</sub>NO | 75.11 | 76.07630 (+H) | 4.58 |
| Glucose | C<sub>6</sub>H<sub>12</sub>O<sub>6</sub> | 180.156 | 198.09677 (+NH<sub>4</sub>) | 3.78 |
| GDL | C<sub>6</sub>H<sub>10</sub>O<sub>6</sub> | 178.14 | 177.03991 (-H) | 2.17 |

**Table S3-S6:** Quantification of selected ions by LC/MS. Excel sheet available online. Contains quantitation results and standard curves (Table S3) of TMA, TMAO, glucose and GDL used in Figure 2 (Table S4) and Figure 4 (Table S5-S6).

**Table S7:** Description of smell criteria for dry hydrolysates, employed by trained sensory panel.

| Parameter | Description |
| --- | --- |
| **Total smell intensity** | Intensity of all smells in the sample |
| **Trimethylamine smell** | The smell of trimethylamine (TMA) |
| **Sweet smell** | Related to a sweet smell |
| **Sour/fermented smell** | A fermented sour smell, spoiled (the smell of a sour dishrag) |
| **Mineral smell** | Related to smells of plaster, lime, chalk and dryness |
| **Seaweed smell** | Related to fresh and dried seaweed and greens, green tea |
| **Feed smell** | Related to the smell of fish feed |
| **Oxidized smell** | Related to an oxidized smell which reminds you of dust, drawers and cardboard |
| **Rancid smell** | The intensity of all rancid smells (grass, hay, candle wax, paint, tallow, soap) |

**Table S8:** Hedonic scale description, used in the consumer panel.

| Score | Acceptability description |
| --- | --- |
| **1** | Dislike very much |
| **2** | Dislike moderately |
| **3** | Dislike slightly |
| **4** | Neither like nor dislike |
| **5** | Like slightly |
| **6** | Like moderately |
| **7** | Like very much |

**Table S9:** Description of odor parameters used in the consumer panel.

| Odor parameter | Description |
| --- | --- |
| **Fishy smell** | Degree to which a fishy odor is perceived |
| **Smell intensity** | Overall intensity of the sample's odor |
| **Smell freshness** | Perception of how fresh the smell is (pleasant, non-stale) |
| **Sulfur-like smell** | Perception of any sulfur-related notes (rotten egg, pungent) |
| **Ammonia-like smell** | Detection of ammonia or any sharp chemical odors |

